# Thermal and alkaline pre-treatments of inoculum halt methanogenesis and enables cheese whey valorization by batch acidogenic fermentation

**DOI:** 10.1101/2023.02.22.529517

**Authors:** Maria Paula Giulianetti de Almeida, Camille Mondini, Guillaume Bruant, Julien Tremblay, David G. Weissbrodt, Gustavo Mockaitis

## Abstract

Carboxylates like volatile fatty acids (VFAs) can be produced by acidogenic fermentation (AF) of dairy wastes like cheese whey, a massive residue produced at 160.67 million m^3^ of which 42% are not valorized and impact the environment. In mixed-culture fermentations, selection pressures are needed to favor AF and halt methanogenesis. Inoculum pre-treatment was studied here as selective pressure for AF demineralized cheese whey in batch processes. Alkaline (NaOH, pH 8.0, 6 h) and thermal (90°C for 5 min, ice-bath until 23°C) pre-treatments, were tested together with batch operations run at initial pH 7.0 and 9.0, food-to-microorganism (F/M) ratios of 0.5 to 4.0 g COD g^-1^ VS, and under pressurized and non-pressurized headspace, in experiments duplicated in two institutes. Acetic acid was highly produced (1.36 and 1.40 g COD_AcOH_ L^-1^) at the expense of methanogenesis by combining a thermal pre-treatment of inoculum with a non-pressurized batch operation started at pH 9.0. Microbial communities comprised of VFAs and alcohol producers, such as *Clostridium*, *Fonticella*, and *Intestinimonas*, and fermenters such as *Longilinea* and *Leptolinea*. Communities also presented the lipid-accumulating and bulk and foaming *Candidatus Microthrix* and the metanogenic *Methanosaeta* regardless of no methane production. An F/M ratio of 0.5 g COD g^-1^ VS led to the best VFA production of 1,769.38 mg L^-1^. Overall, inoculum thermal pre-treatment, initial pH 9.0, and non-pressurized headspace acted as a selective pressure for halting methanogen and producing VFAs, valorizing cheese whey via batch acidogenic fermentation.

## 1 Introduction

Anaerobic digestion (AD) has been used from the 19^th^ century onwards to obtain biogas as an energy carrier (Kigozi et al., 2014). Since then, AD processes are being constantly improved (Abbasi et al., 2012; van Lier et al., 2001) to process various waste streams (Ostrem et al., 2004) and address energy production, environmental burdens, and circular economy. The primary goal remains to reduce organic matter and remove nutrients from agro-industrial, food, and municipal solid wastes (Chen et al., 2008; Goud and Mohan, 2012; Wonglertarak and Wichitsathian, 2014).

Cheese whey (CW) is a by-product of the dairy industry with a high organic load (i.e., 50 to 80 g O_2_ L^-1^, in terms of chemical oxygen demand (COD))(Saddoud et al., 2007). It has an annual production of 160.67 million m^3^ where 58% of the production is absorbed by various industries (e.g., food, nutrition, cosmetics, and pharmaceutical). However, a staggering amount of 66.5 million m^3^ year ^-1^ of CW (Food and Agriculture Organization of the United Nations, n.d.; Tsakali et al., 2010; USDA - Foreign Agricultural Service, 2022) is currently transformed into low added-value products such as animal feed, fertilizer, or being discharged into water bodies leading to eutrophication processes (Smithers, 2008; Tsakali et al., 2010). Hence, cheese whey can be an excellent substrate for AF processes.

Acidogenic fermentation focuses on the valorization of organic matter via the carboxylate platform, by combining the inhibition of methanogenesis and production of VFAs at high yields, which are of great economic interest due to their potential industrial applications (e.g., biofuels, biopolymers, and chemicals) (Angenent et al., 2016; Kleerebezem et al., 2015; Mulders et al., 2020; Rombouts et al., 2019).

Selective pressures (e.g., pH, physicochemical pre-treatments, reactor headspace pressure, and F/M ratio) can dictate microbial diversity and dynamics, interactions, energy requirements, and preferred metabolisms (Pretorius, 1987). Most studies in AD and AF processes focus on the pre-treatment of the substrate and waste-activated sludge (Sarkar et al., 2021). The present study focused on various selective pressure mechanisms for enhancing VFA production and halting methanogenesis in AF processes. We also investigated the effects of alkaline and thermal inoculum pre-treatments, variations in the initial pH in headspace pressure, and different F/M ratios on the acidogenic fermentation of cheese whey.

Alkaline pre-treatments have been shown to improve AD processes by increasing sludge solubilization and enhancing methane production (Chen et al., 2007; Li et al., 2012; López-Torres and Llórens, 2016; Navia and Vidal, 2002; Wonglertarak and Wichitsathian, 2014), while thermal pre-treatments have been used to inhibit methanogenesis (Alibardi et al., 2012; Ramos-Suarez et al., 2021; Sarkar et al., 2017).

However, thermal pre-treatments, that can be performed at mesophilic or hyperthermophilic temperatures (i.e., 30°C to 180°C) are time-consuming (up to several hours), which would increase AF processes’ overall costs (Corti and Lombardi, 2007). Contradictory results have also been observed regarding the effects of such pre-treatments on methanogenesis, with an inhibition observed after alkaline pre-treatment (Yuan et al., 2006) and an enhancement observed after thermal pre-treatment (Zhang et al., 2015). The lack of consensus on the best conditions for inoculum pre-treatment and their influence on AF processes still needs to be elucidated. To date, studies on thermal pre-treatments are mostly performed on waste-activated sludge. Their use as a selective pressure mechanism for VFA production remains scarce.

Variations in the initial pH can positively influence VFA production (Dareioti et al., 2014). In addition, uncontrolled pH approaches can reduce AF process costs since there is no further need for chemical utilization for stabilizing pH (Sarkar et al., 2021).

Headspace gas composition and pressure play a role in product formation and metabolic pathways preferentially used. However, most studies still focus on hydrogen production (Darvekar et al., 2019). According to Zhou (2018) and Sarkar (2017), low hydrogen pressure favors VFA formation. Finally, the F/M ratio, which is inoculum and substrate-dependent, is a parameter that impacts acidogenesis, with lower F/M ratios being beneficial to VFAs production (Pang et al., 2019; Shah et al., 2015).

In this work, we aimed at (*i*) identifying how abiotic factors (i.e., pH, inoculum pre-treatment, headspace pressure, and F/M ratio) influence the products spectra of cheese whey via AF, (*ii*) validating thermal pre-treatment efficiency for halting methanogenesis, and (*iii*) identifying the parameters that increase acetate level of production for other biological processes (e.g., microalgal photoorganoheterotrophic biomass production).

## 2 Materials & Methods

Figure 1 depicts the overall methodology, involving experiments performed at the University of Campinas (UNICAMP, Brazil) and reproduced at Delft University of Technology (TU Delft, The Netherlands). Substrate and inoculum preparation and inoculum pre-treatment are common to all experiments apart from a few modifications described hereafter.

**Figure 1.**
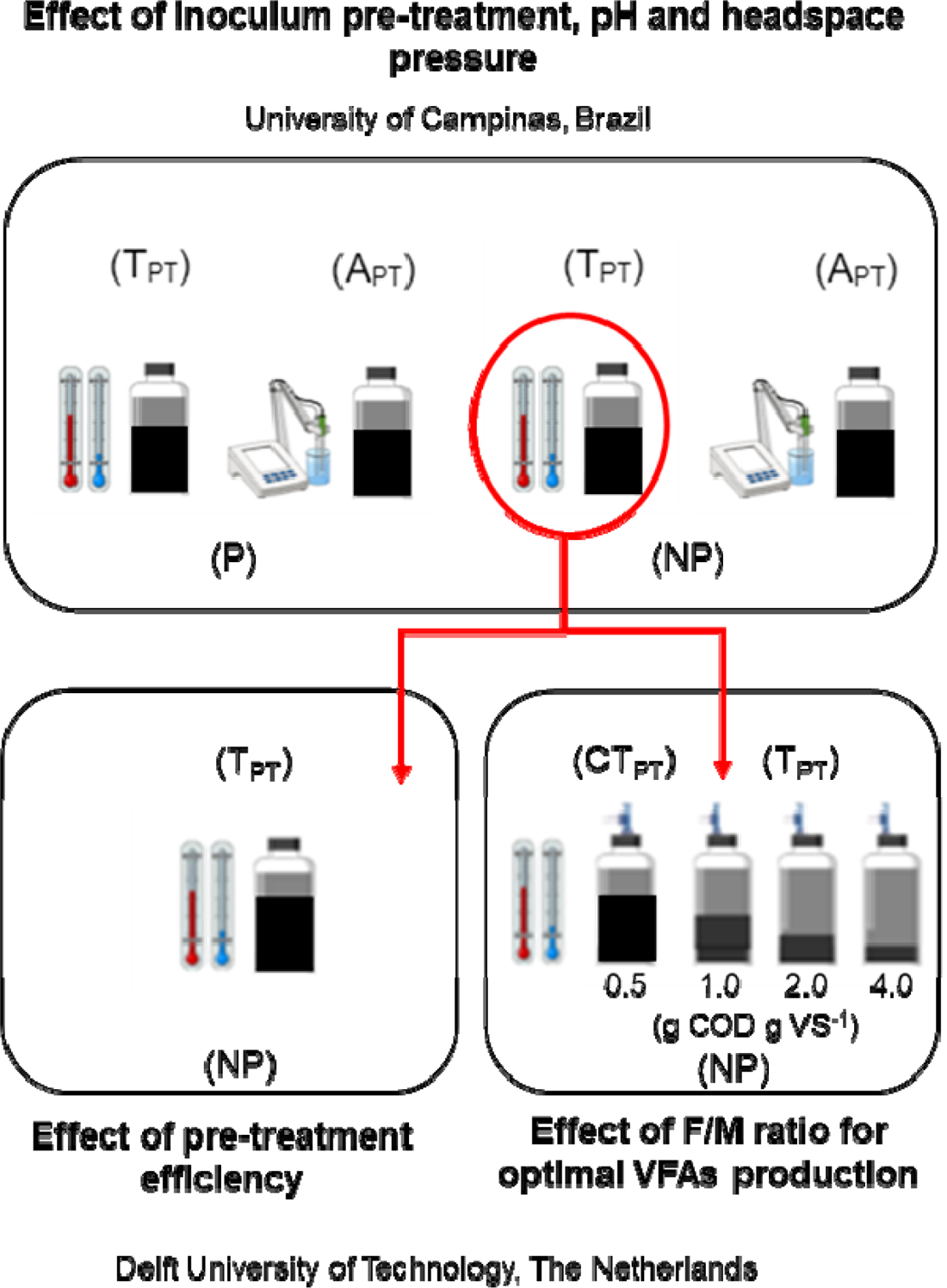
Overall experiment methodology. The inoculum went through alkaline (A_PT_) and thermal (T_PT_) pre-treatments. The initial pH at the start of the digestions was set at 7.0 and 9.0 and headspace assays were carried in pressurized (P) and non-pressurized (NP) batches. Analyses encompassed gas and VFA profiles, total organic carbon (TOC), total nitrogen (TN), COD, solid series, carbohydrates, and amplicon sequencing (16S rRNA gene).

### 2.1 Demineralized cheese whey media

The chosen carbon source for all experiments consisted of 40% demineralized whey powder (WPC40), which was the closest substrate to raw cheese whey. The substrate medium for experiments held at UNICAMP was composed of (per L): WPC40 (Pic-Nic, Brazil) (4 g) (Mockaitis et al., 2006) and the following reagents: NaHCO_3_ (1 g), NaCl (250 μg), MgCl_2_.6H_2_O (7 μg) and CaCl_2_·2H_2_O (9.5 μg) (Torres, 1992). The medium was prepared fresh before experiments and its initial pH was approximately 7.0. No pH corrections were made after medium preparation. The sludge used as inoculum was obtained from a UASB reactor from a poultry abattoir in Brazil (25°05’10.1” S 47°58’49.7” W). National System for the Management of Genetic Heritage and Associated Traditional Knowledge (SisGen) register number is AE43468.

For the experiments held at TU Delft, the medium was composed of (per L): WPC40 (Subo International, The Netherlands) (4 g) amended with 0.3 mL of a micronutrients solution per g of substrate COD. The micronutrients solution was composed of FeCl_3_·6H_2_O (2 g L^-1^), CoCl_2_·6H_2_O (2 g L^-1^), MnCl_2_·4H_2_O (0.5 g L^-1^), CuCl_2_·2H_2_O (32 mg L^-1^), ZnCl_2_ (50 mg L^-1^), HBO3 (50 mg L^-1^), (NH_4_)6Mo_7_O_2_·4H_2_O (90 mg L^-1^), Na_2_SeO_3_·5H_2_0 (100 mg L^-1^), NiCl_2_·6H_2_O (6 mg L^-1^), EDTA (1 g L^-1^), HCl 36% (1 mL L^-1^), Resazurine (0.5 g L^-1^) and yeast extract (2 g L^-1^). Sludge used as inoculum was obtained from an anaerobic digester at WWTP Harnashpolder in The Netherlands (52°00’48.8“N 4°19’02.5”E). It is valid to notice that 1 g L^-1^ of WPC40 corresponds to 1 g O_2_ L^-1^ in COD terms.

### 2.2 Mechanical homogenization, thermal and alkaline pre-treatments of inocula

The anaerobic granular sludge used as inoculum at UNICAMP was mechanically homogenized by using a blender (Britannia, Brazil) to decrease the size of bioaggregates while increasing their superficial area. The anaerobic granular sludge used as inoculum in TU Delft was used as is.

Thermal pre-treatment, which was performed both at UNICAMP and TU Delft, was adapted from Mockaitis (2020). Briefly, the sludge was heated at 90°C in a water bath for 20 minutes under constant stirring. Heating was then halted by decreasing the sludge temperature to 23°C in an ice bath.

Alkaline pre-treatment consisted in increasing the sludge pH up to 8.0 using 1 M NaOH under continuous mixing, and controlling it for the next 6 hours, with a correction to 8.0 if needed. This pre-treatment was only performed at UNICAMP.

### 2.3 Effect of thermal and alkaline pre-treatments of inocula, initial batch pH, and headspace pressure on acidogenic fermentation

Batch experiments were conducted at UNICAMP. They assessed the effects of three factors evaluated at two levels in WPC40 acidogenic fermentation, namely: (*i*) inoculum pre-treatment (alkaline (A_PT_) and thermal (T_PT_)) to enhance sludge biodegradability and select for alkaline-tolerant acetoclastic microorganisms, and to inactivate methanogenic microorganisms, respectively; (*ii*) variation in the initial pH of digestion process (7.0 and 9.0) to further select microorganisms adaptable to neutral and alkaline pH environments, and (*iii*) gas headspace pressure (pressurized (P) and non-pressurized (NP)) to investigate the influence on AF products spectra. Table 1 shows the two-level factorial experimental design with 3 factors (2^3^ designs) applied to this setup.

**Table 1.**
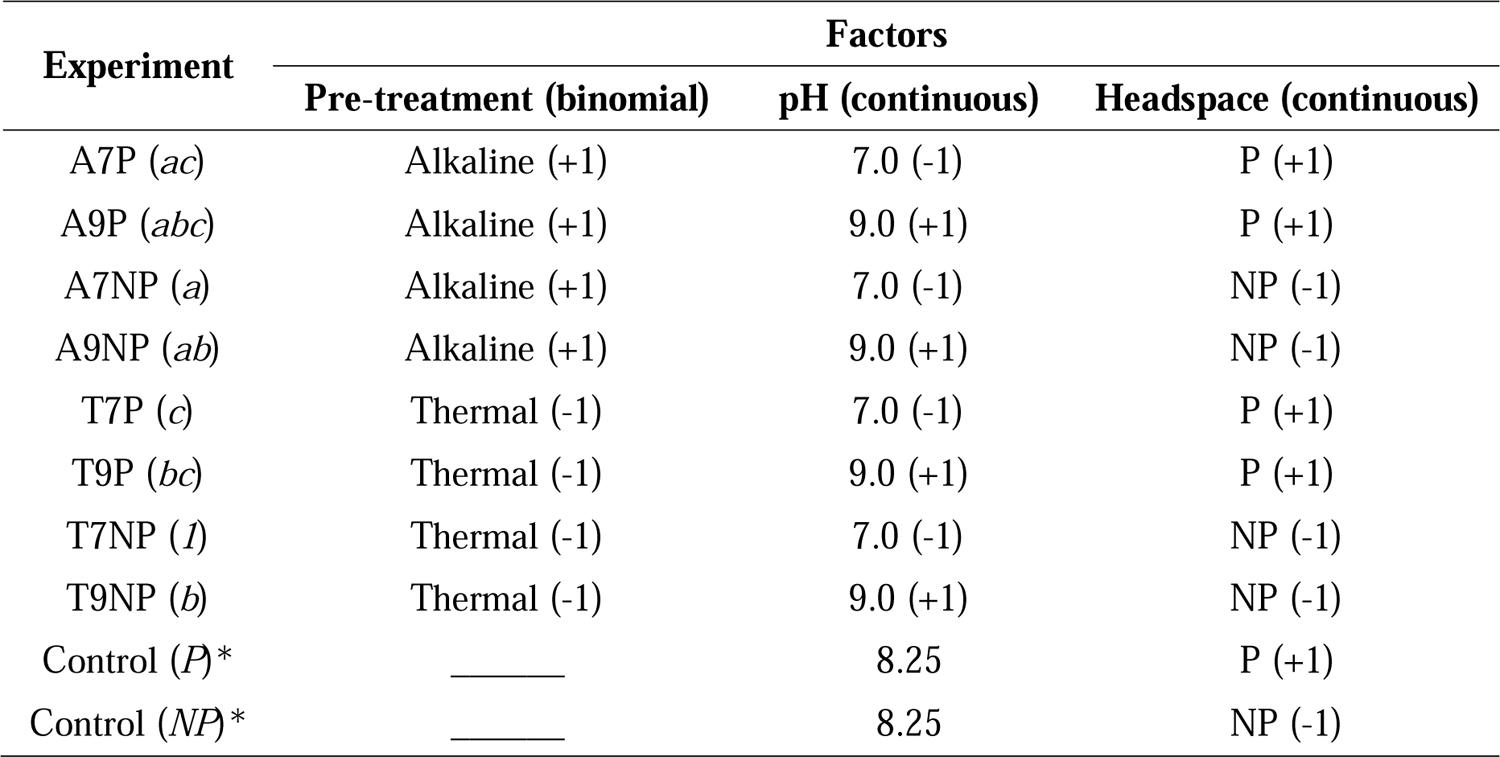
The 23 experimental designs for phase one. The (±1) values indicate the upper and lower levels of the investigated factors. The design aimed at testing the main effects (referred to as a, b and c), the 2-factor interactions effects (ab, ac, and bc), and the 3-factors interaction effect (abc). Two control experiments were included.

Experiments (Figure 1) were performed in 1 L Duran flasks (total volume of 1,130 mL) with an initial working volume of 730 mL and an initial headspace volume of 398 mL. Bottles were inoculated at a concentration of 6.6 g total volatile solids (TVS) L^-1^, as depicted in Mockaitis et al. (2020). Batch reactors were continuously agitated at 50 rpm (Orbital Incubator Marconi – MA420) at a mesophilic temperature of 35°C for 30 days.

#### 2.3.1 Batches headspace pressure assays

The influence of headspace pressure (pressurized and non-pressurized batches) on VFAs production was investigated according to Peixoto et al., (2011). Manometric pressure was measured, then, 3 mL samples of gas were collected with a syringe containing a pressure lock (Thermo Fisher). Each sample corresponded to a batch condition.

In pressurized headspace assays, butyl rubber stoppers of each flask were covered with silicone sealant after manometric measurement and gas sample collection, allowing gas accumulation in the batch headspace.

In non-pressurized experiments, the headspace of each flask was punctured with a needle after pressure manometric reading and gas sampling, allowing the remaining gas to be released until reaching the atmospheric pressure value. After this step, butyl rubber stoppers were also covered with silicone sealant avoiding any gas release into the atmosphere.

Once collected, gas samples were analyzed with a gas chromatograph (GC) equipped with a thermal conductivity detector GC-TCD (Construmac, Brazil) with hydrogen as a carrier gas.

The volumetric production of biogas and its constituents were inferred through equation 1 as a punctual function from sampling timestep *t_0_* = 0 to *t* for non-pressurized assays.

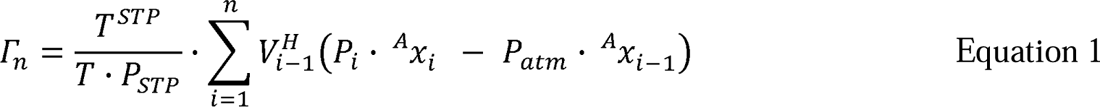

Where *Γ_n_* is the volumetric production of a gas of interest (N_2_, H_2_, CH_4_ or CO_2_) in standard temperature and pressure (STP) equivalents (L), at a given time *n*; *P_i_* is the measured pressure at sampling time (atm); *V^H^_i-1_* is the headspace volume before sampling (L); *T* is system’s temperature (K), *T^STP^* is the standard temperature (273 K), *P^STP^* is standard pressure (1 atm), *^A^X_i_* is the molar fraction of the gas of interest at sampling time and *^A^X_i-1_* is the molar fraction of the gas of interest before sampling.

#### 2.3.2 VFAs measurements

VFAs, alcohols and carbohydrates were quantified by high performance liquid chromatography (HPLC) based on Penteado et al.,(2012). The VFAs and alcohol profile consisted of the following: lactic acid, formic acid, acetic acid, propionic acid, butyric acid, iso-butyric acid, iso-valeric acid and ethanol.

The chromatograph was equipped with two LC-20AD pumps, one DGU-20A3R degasser, a SIL-20AHT autosampler, a CTO-20A column oven, a SPD-20 UV detector with readings at 210 nm (Shimadzu, Japan), an Aminex HPX-87H 300 x 7.8 mm column (BioRad, USA), a RID-10A index refraction detector and a CBM-20A controller (Shimadzu, Japan).

Two mL of mixed liquors were centrifuged at 16,025 xg for 5 min.

Forty μL of H_2_SO_4_ (2M) was added to 1 mL of the supernatant to acidify the sample for ideal analyte separation. The samples were then filtered through regenerated cellulose (RC) syringe filters (RC 0.20 μm) (GVS, Italy), and transferred to 1.5 mL pre-washed vials (H_2_SO_4_ at 2 mol L^-1^ to avoid any contamination) before HPLC analyses. The HPLC run time was 60 minutes per sample with a constant column oven temperature of 43°C and 0.005 M H_2_SO_4_ mobile phase at a flux rate of 0.5 mL min ^-1^.

### 2.4 Inoculum thermal pre-treatment and non-pressurized headspace efficiency

Inoculum thermal pre-treatment and non-pressurized headspace assays were reproduced to confirm the efficiency of such imposed conditions in halting methanogenesis while enhancing VFAs production. Each factor was tested at one level, based on the effects identified at UNICAMP (section 2.3): inoculum thermal pre-treatment, initial pH of 9.0, and non-pressurized headspace. A batch with no inoculum pre-treatment, initial pH of 9.0, and non-pressurized headspace acted as a control. The experimental design with three factors applied to this set-up is shown in Table 2.

**Table 2.** Experimental design for the F/M ratio experiments. All experiments were done in duplicate.

| Experiment | Factors |  |  |
| --- | --- | --- | --- |
|  | Pre-treatment | F/M ratio (continuous) | Headspace (continuous) |
| FM0.5 | Thermal | 0.5 | NP |
| FM0.5 | Thermal | 0.5 | NP |
| FM1.0 | Thermal | 1.0 | NP |
| FM1.0 | Thermal | 1.0 | NP |
| FM2.0 | Thermal | 2.0 | NP |
| FM2.0 | Thermal | 2.0 | NP |
| FM4.0 | Thermal | 4.0 | NP |
| FM4.0 | Thermal | 4.0 | NP |

Similar to the first experiments conducted, the initial working volume of the 1 L Duran flasks was 750 mL with an initial headspace volume of 398 mL, for a total working volume of 1,148 mL. The inoculum concentration was identical at 6.6 g TVS L^-1^ as depicted in Mockaitis et al., (2020). Batch reactors were incubated for 10 days at a mesophilic temperature of 35°C in a Certomat^®^ BS1 incubator (Sartorius Stedim Biotech, Germany) under continuous agitation (145 rpm). The headspace was released with the aid of Cole-Palmer stopcocks with Luer connection, a 1-way male lock (Cole-Parmer, USA). Experiments were performed in triplicate and lasted 12 days.

#### 2.4.1 VFAs measurements

VFAs and carbohydrates were quantified by HPLC. One mL of mixed liquors was centrifuged at 5,073 x g for 5 minutes and supernatants were filtered using a 0.45 μm syringe filters. Filtered samples were then transferred to 300 µL Waters total recovery vials, from which 10 µL were injected with a 2707 Waters HPLC autosampler at 15°C The chromatograph was equipped with a Waters M515 HPLC (Waters, USA) pump, a Waters 2414 refractive index (RI) detector, with 1,024 of sensitivity and a Waters 2489 UV/Visible detector at 210 nm. The column was a BioRad HPX-87H (300 x 7.8 mm) with a BioRad Cation-H refill cartridge (30 x 4.6 mm) guard column (BioRad, USA) and the column oven was built in-house. The flow rate of the pump was 0.6 mL min^-1^ and the temperature of the column was set at 59°C. The RI detector was operated at 30 °C. The mobile phase was 1.5 mmol L^-1^ phosphoric acid diluted in ultrapure water (MilliQ, Merck Millipore). The VFAs measured were acetic acid, butyric acid, formic acid, caproic acid, propionic acid, valeric acid, iso-butyric acid, iso-caproic acid, and iso-valeric acid.

### 2.5 Ideal F/M ratio for optimal VFAs production with thermally pre-treated inocula

F/M ratio is a measurement used to determine the amount of substrate needed for the number of microorganisms present in a system. It is an important parameter to evaluate in an approach aiming at maximizing VFA production. To determine the best FM ratio for VFA production when working with a thermal pre-treated sludge at non-pressurized headspace, four different F/M ratios were tested: 0.5, 1.0, 2.0, and 4.0,g_COD_ g_VS_ ^-1^. These ratios were obtained by dividing the COD of the substrate by the VS of the sludge. The initial pH of all experiments was 9.0. In section 2.4, both control and thermal batches presented an F/M ratio of 0.5 g _COD_ g _VS_ ^-1^. g _COD_ g _VS_ ^-1^ acted as control. Assays were conducted in duplicates for 14 days. The experimental design is depicted in Table 2.

#### 2.5.1 Biogas measurements

The presence or absence of CH_4_ in the gas was detected by injecting 10 mL samples of gas from the headspace of the bottles in a GC (Agilenttech 7890 A, Agilent Technologies Inc., USA) equipped with an HP-PLOT Moleseive GC column (Agilent 19095P-MS6, Agilent Technologies Inc., USA) of 60 m x 0.53 mm x 200 µm and a thermal conductivity detector (TCD). The carrier gas was helium (14.8 psi, 23 mL min^-1^) and the operating temperature was 200 °C.

### 2.6 Physicochemical analyses

Mixed liquors and gas phases of the experimental flasks were sampled for physicochemical analyses in both original and replicated experiments. For the experiments held at UNICAMP, every 15 days, 150 mL of mixed liquors were collected and used in the following analyses and respective code methods: solid series (2540 B-F; 50 mL), COD (5220 B; 1 mL), sulfate (4500-SO_4_^2−^ E; 1 mL), sulfide (4500-S^2^ D, 1 mL), total organic carbon (5310-TOC; 15 mL) (APHA, 2005) and total nitrogen (TN, ASTM D8083; 15 mL) (ASTM International, 2016). Additional analyses were performed consisting of pH measurements (4500 H) (Federation 2005), headspace assays (CH_4_, N_2,_ and CO_2_, 3 mL of gas phase), HPLC (2 mL) for VFAs, sugars and alcohols characterization, total carbohydrates measurements (Dubois et al., 1956) (1 mL), total alkalinity and VFAs measurements (Dilallo and Albertison, 1961; Ripley, L.E., Boyle, W.C. and Converse, 1986) (60 mL). The remainder sludge was discharged. On alternate days, these analyses were also performed.

For the replicated experiments performed at TU Delft, 100 mL of mixed liquors were collected on the first and last days of the process and were used in the following analyses in the same volume as mentioned above: solid series, COD, and TOC/TN. Other analyses were also performed, consisting of pH measurements, biogas composition determination, HPLC for VFAs and sugars characterization, and total carbohydrates measurements. In addition, on alternate days 12 mL of mixed liquors were sampled for pH, HPLC, COD, TOC, and TN measurements.

### 2.7 16S rRNA gene amplicon sequencing

#### 2.7.1 Sequencing experiments

Microbial community analyses were performed through 16S rRNA gene amplicon sequencing on (*i*) the initial and final samples from inoculum thermal pre-treatment efficiency assays and (*ii*) the final samples of each F/M ratio experiment.

0.5 mL samples of mixed liquors were collected from each condition in 1.5 mL Eppendorf tubes completed with demineralized water for initial washing. Samples were centrifuged at 10,000 x g, 4 °C for 3 minutes, and the supernatant was discarded. DNA was extracted using the Qiagen DNeasy® UltraClean® Microbial Kit (Qiagen, Germany) according to the manufacturer’s instructions. The quality and concentrations of the DNA extracts were assessed using a Qubit fluorometer and the Qubit dsDNA HS assay kit (Thermo Fischer Scientific, USA) according to the manufacturer’s instructions.

For each sample, a minimum of 20 μL of DNA extracts with concentrations ranging from 13.9 and 95 ng μL^-1^ were sent to Novogene (Beijing, China) for 16S rRNA gene amplicon sequencing. All sequencing experiments (i.e 16S rRNA sequence amplifications, libraries preparations, and sequencing runs) were performed at Novogene, according to their protocols. The targeted hypervariable regions V3-V4 of the 16S rRNA gene were amplified by PCR with the pair of barcoded primers 341F (CCTAYGGGRBGCASCAG) and 806R (GGACTACNNGGGTATCTAAT) (Yu et al., 2005). This set of primers is known to cover both bacteria and archaea with high specificity, except for *Planctomycetes* (Weissbrodt et al., 2020).

#### 2.7.2 Sequences analyses

All sequencing data received from Novogene were analyzed using AmpliconTagger (Julien Tremblay, 2019). Briefly, raw reads were scanned for sequencing adapters and PhiX spike in sequences. The remaining paired-end reads were processed to remove primer sequences (pTrimmer v1.3.3 – (Zhang et al., 2019) and discard reads having average quality Phred score lower than 20. The remaining sequences were processed for generating Amplicon Sequence Variants (ASVs) (DADA2 v1.12.1) (Callahan 2016) using the following key parameters: filterAndTrim (maxEE =2; truncQ = 0; maxN = 0; minQ = 0) and learnErrors (nbases=1e8) functions for both forward and reverse filtered reads. Reads were then merged using the mergePairs (minOverlap = 10; maxMismatch = 0) function. Chimeras were removed with DADA2’s internal removeBimeraDeNovo (method = ‘consensus’) method followed by UCHIME reference (Rognes et al., 2016). ASVs were assigned a taxonomic lineage with the RDP classifier (Wang et al., 2007) using training sets containing the complete Silva release 138 database (Quast et al., 2013) supplemented with a customized set of mitochondria and plastid sequences. The RDP classifier gave a score of (0 to 1) to each taxonomic depth of each ASV. Any taxonomic depth having a score of ≥ 0.5 was kept reconstructing the final lineage. Taxonomic lineages were combined with the cluster abundance matrix obtained above to generate a raw ASV table from which the bacterial or fungi ASV tables were generated. Five hundred 1,000 reads rarefactions were then performed on these ASV tables and the average number of reads of each ASV of each sample was computed to obtain consensus rarefied ASV tables. Alpha diversity metrics (RTK v 0.93.2 – ((Saary et al., 2017) and taxonomic summaries microbiomeutils v 0.9.3 – (Julien Tremblay, 2020)) were then computed using the consensus rarefied ASV tables. Figures were generated in R using the ggplot2 package.

## 3 Results and Discussion

### 3.1 Effects of inoculum thermal and alkaline pre-treatments and initial pH on the anaerobic digestion process

#### 3.1.1 Inoculum thermal pre-treatment efficiently halted methanogenesis

As demonstrated in Figure 2, inoculum thermal pre-treatment successfully inhibited methanogenesis in all tested conditions (i.e., pressurized and non-pressurized headspace assays with initial pH of 7.0 and 9.0).

**Figure 2.**
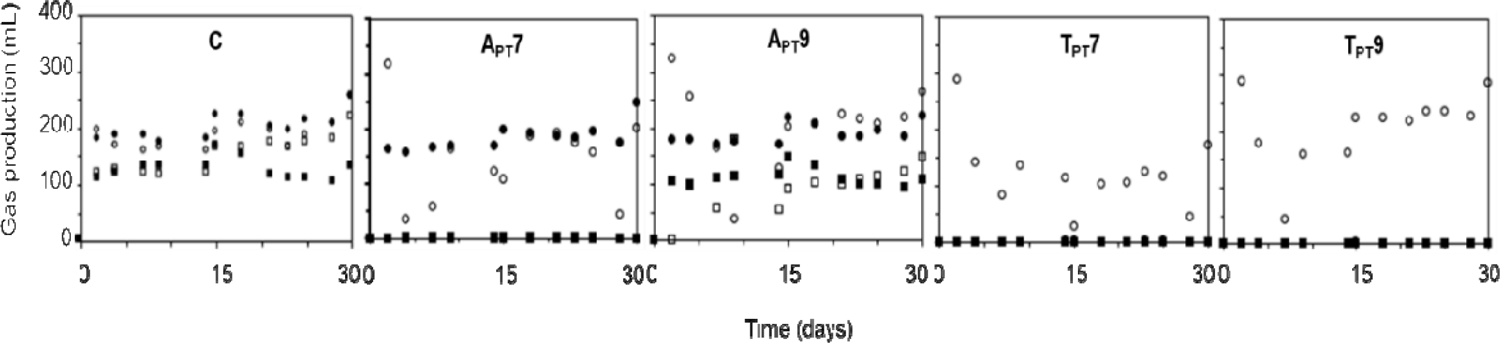
Methane and carbon dioxide gas production profiles for control (C), inoculum alkaline (A_PT_), and thermal (T_PT_) pre-treatment with initial pH of 7.0 and 9.0 during 30 days. Non-pressurized (NP) and Pressurized (P) headspace assays are also represented. In the figure: CH_4_NP,

Inoculum alkaline pre-treatment halted methanogenesis in both pressurized and non-pressurized assays but only with initial pH 7.0, presenting negligible to no levels of total methane production of 0.0 and 0.54 mL, respectively. Initial pH 9.0 presented a total methane production of about 52.29 mL for pressurized assays and 0.67 mL for non-pressurized ones.

As a strategy to maximize VFA production, inoculum thermal pre-treatment appears to be the best selective pressure for impairing methanogens in batch processes. It is of utmost importance to confirm whether this selective pressure is also successful to halt methanogenesis considering different types of inoculum (e.g., waste activated sludge and secondary sludge), reactors (e.g., sequence-batch reactors, and continuous-stirred tank reactors) and configurations (e.g., organic loading rate, hydraulic retention time, coupled reactors). If so, then inoculum thermal pre-treatment can be implemented in full-scale AF processes, decreasing their overall costs (Corti and Lombardi, 2007).

Some studies have used alkaline pre-treatment of inoculum to increase the biodegradability of waste-activated sludge (Wonglertarak and Wichitsathian, 2014), and to stabilize the sludge originated from aerobic processes while treating the mineralization of its remaining organic compounds (Navia and Vidal, 2002). Sludge alkaline pre-treatment increased the COD: COD_Total_ ratio (Navia and Vidal, 2002) VS while improving volatile solids (VS) reduction (Appels et al., 2011; Lin et al., 1997). Hence, their methane production would also increase.

Noteworthily, the methane production levels observed in our experiments over 30 days (i.e., A_PT_7NP, 0.54 mL; A_PT_7P, 0.00 mL; A_PT_9NP 0.67 mL, and A_PT_9P, 52.29 mL), did not agree with the findings of Wonglertarak and Wichitsathian (2014). Their methane production was about 2, 526.60 mL in a thermophilic reactor with alkaline pre-treated sludge and 2,173.8 mL in a mesophilic reactor with alkaline pre-treated sludge. Our findings also contradicted the ones found by Lin et al. (1997) which was an average CH_4_ production rate of 222.25 L m^3^ d^-1^.

This could be explained using different methodologies in each work such as maintaining the reactor with a constant alkaline pH after treating the sludge, different reactors configurations (e.g., single-stage high-rate reactor and continuous stirred tank reactor and batch reactor), hydraulic retention times, organic loading rates as well as the type of inoculum used (i.e., WAS and UASB granular sludge).

For instance, we used 1 M NaOH to treat the inoculum, correcting the pH to 8.0 hourly for 6 hours. However, after pre-treating the inoculum, batch reactors were assembled with initial pH of 7.0 and 9.0, with the final pH from both pressurized and non-pressurized headspaces, ranging from 4.9 and 5.1, respectively. Lin et al. (1997), treated secondary sludge with NaOH ranging from 0.02 to 0.2 M, which pH increased from 7.21 to 13.2 in 24 h hydrolysis and 0.2 M NaOH. Alternatively, Wonglertarak and Wichitsathian’s (2014) alkaline inoculum pre-treatment consisted of treating waste-activated sludge with pH varying from 8.0 to14.0 and varying NaOH concentration from 8 to 31,000 g m^-3^ wet sludge.

Another result of interest regarding inoculum alkaline pre-treatment was the correlation between acetate and methane production. While there was no methane production in inoculum alkaline treatment with initial pH 7.0 in both non-pressurized and pressurized headspace batches, acetate production differed 185.6% from non-pressurized headspace assay to pressurized headspace (i.e., A_PT_7NP 1,056.04 mg COD_AcOH_ L^-1^ and A_PT_7P 369.82 mg COD_AcOH_ L^-1^). So, headspace pressure can favor acetate production in acetoclastic methanogens.

The detected levels of CO_2_ in both A_PT_7NP and A_PT_7P (i.e., 1.84 mL and 60.78 mL, respectively) were low as was the average of the final pH for both conditions (i.e., 4.8). Hence, instead of methanogenesis being inhibited due to disturbance in the buffer system (Casallas-Ojeda et al., 2020), we can infer that most CO_2_ was dissolved in the liquid phase, decreasing the pH (Deublein and Steinhauser, 2011). Figure 2 shows the methane and carbon dioxide production in alkaline and thermal batches with both pressurized and non-pressurized headspaces.

#### 3.1.2 Initial pH, headspace pressure, and inoculum pre-treatments influenced VFAs product spectra

Cheese whey has a natural tendency to acidify and since pH was not controlled during the experiments (pressurized and non-pressurized batches displayed an average pH of 4.96 and 5.26, respectively), it is likely that hydrolysis was not a limiting step in the AF process.

An additional hypothesis to justify the different production of VFAs and alcohol in all batches was that together with uncontrolled pH, the headspace internal pressure variation from pressurized and non-pressurized batches played a key role on the spectra and quantity of VFAs produced. In pressurized batches, the pressure would contribute to the gas phase being accumulated within the liquid phase, which influenced both gas and VFA spectra. In non-pressurized batches, due to headspace release, gas and VFAs production restarted daily, after each sampling.

Overall, non-pressurized batches produced more VFAs than pressurized ones. Special highlights can be given to A_PT_9 NP (21,360.54 mg COD_compound_ L^-1^), T_PT_9 NP (4,108.71 mg COD_compound_ L^-1^), and T_PT_7 NP (3,414.15 mg COD_compound_ L^-1^), when compared to their counterparts (A_PT_9 P 1,940.14 mg COD_compound_ L^-1^, T_PT_9 P 2,151.99 mg COD_compound_ L^-1^ and T_PT_7 P 2,140.62 mg COD_compound_ L^-1^). Both pressurized and non-pressurized controls and A_PT_7 did not present a substantial difference with an increase of approximately 2.9% in VFAs produced in non-pressurized batches compared to pressurized batches as seen in table 3.

**Table 3.**
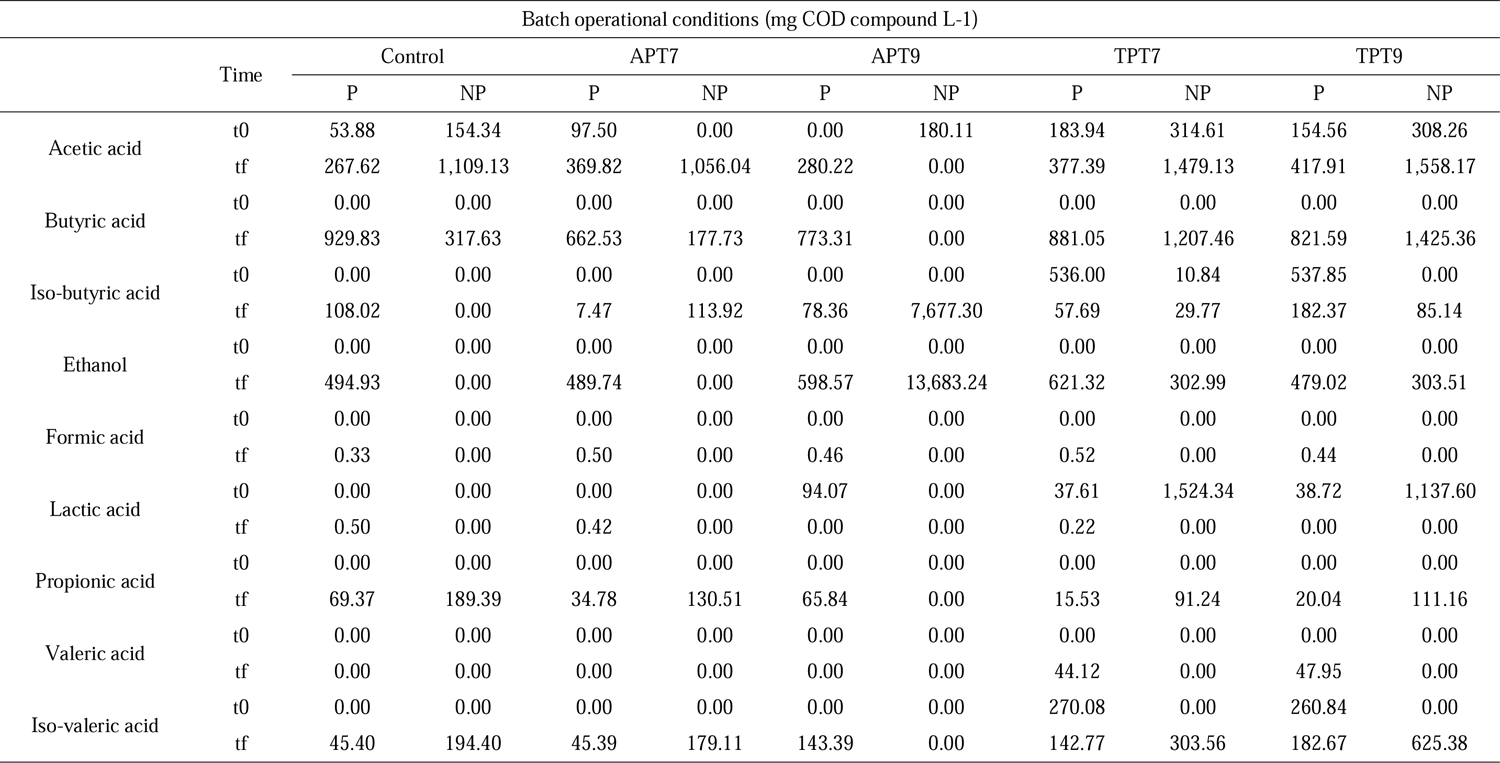
VFAs and alcohols product spectra of pressurized (P) and non-pressurized (NP) headspace batches with alkaline (A_PT_) and thermal (T_PT_) pre-treatments and initial pH of 7.0 and 9.0. Experiments lasted for 30 days.

| Batch operational conditions (mg COD compound L-1) |  |  |  |  |  |  |  |  |  |  |  |
| --- | --- | --- | --- | --- | --- | --- | --- | --- | --- | --- | --- |
|  | Time | Control |  | APT7 |  | APT9 |  | TPT7 |  | TPT9 |  |
|  |  | P | NP | P | NP | P | NP | P | NP | P | NP |
| Acetic acid | t0 | 53.88 | 154.34 | 97.50 | 0.00 | 0.00 | 180.11 | 183.94 | 314.61 | 154.56 | 308.26 |
|  | tf | 267.62 | 1,109.13 | 369.82 | 1,056.04 | 280.22 | 0.00 | 377.39 | 1,479.13 | 417.91 | 1,558.17 |
| Butyric acid | t0 | 0.00 | 0.00 | 0.00 | 0.00 | 0.00 | 0.00 | 0.00 | 0.00 | 0.00 | 0.00 |
|  | tf | 929.83 | 317.63 | 662.53 | 177.73 | 773.31 | 0.00 | 881.05 | 1,207.46 | 821.59 | 1,425.36 |
| Iso-butyric acid | t0 | 0.00 | 0.00 | 0.00 | 0.00 | 0.00 | 0.00 | 536.00 | 10.84 | 537.85 | 0.00 |
|  | tf | 108.02 | 0.00 | 7.47 | 113.92 | 78.36 | 7,677.30 | 57.69 | 29.77 | 182.37 | 85.14 |
| Ethanol | t0 | 0.00 | 0.00 | 0.00 | 0.00 | 0.00 | 0.00 | 0.00 | 0.00 | 0.00 | 0.00 |
|  | tf | 494.93 | 0.00 | 489.74 | 0.00 | 598.57 | 13,683.24 | 621.32 | 302.99 | 479.02 | 303.51 |
| Formic acid | t0 | 0.00 | 0.00 | 0.00 | 0.00 | 0.00 | 0.00 | 0.00 | 0.00 | 0.00 | 0.00 |
|  | tf | 0.33 | 0.00 | 0.50 | 0.00 | 0.46 | 0.00 | 0.52 | 0.00 | 0.44 | 0.00 |
| Lactic acid | t0 | 0.00 | 0.00 | 0.00 | 0.00 | 94.07 | 0.00 | 37.61 | 1,524.34 | 38.72 | 1,137.60 |
|  | tf | 0.50 | 0.00 | 0.42 | 0.00 | 0.00 | 0.00 | 0.22 | 0.00 | 0.00 | 0.00 |
| Propionic acid | t0 | 0.00 | 0.00 | 0.00 | 0.00 | 0.00 | 0.00 | 0.00 | 0.00 | 0.00 | 0.00 |
|  | tf | 69.37 | 189.39 | 34.78 | 130.51 | 65.84 | 0.00 | 15.53 | 91.24 | 20.04 | 111.16 |
| Valeric acid | t0 | 0.00 | 0.00 | 0.00 | 0.00 | 0.00 | 0.00 | 0.00 | 0.00 | 0.00 | 0.00 |
|  | tf | 0.00 | 0.00 | 0.00 | 0.00 | 0.00 | 0.00 | 44.12 | 0.00 | 47.95 | 0.00 |
| Iso-valeric acid | t0 | 0.00 | 0.00 | 0.00 | 0.00 | 0.00 | 0.00 | 270.08 | 0.00 | 260.84 | 0.00 |
|  | tf | 45.40 | 194.40 | 45.39 | 179.11 | 143.39 | 0.00 | 142.77 | 303.56 | 182.67 | 625.38 |

It is important to stress that although A_PT_9 NP, T_PT_9 NP, and T_PT_7 NP produced greater quantities of metabolites, they did not produce a great variety of VFAs and alcohol. A_PT_9 NP produced 7,677.30 mg COD_IBA_ L^-1^ of iso-butyric acid and 13,683.24 mg COD_EtOH_ L^-1^ of ethanol. These metabolites production together with the absence of acetic acid production are a clear result of a metabolic shift that was favored due to the imposed selective pressures in this batch (i.e., alkaline inoculum pre-treatment, initial pH of 9.0, and non-pressurized headspace).

This was in opposition to all other conditions tested. It is likely that in such operational conditions, acetate served as the substrate for fermentative bacteria to form iso-butyric acid and ethanol, as already described by Thatikayala et al. (2021) and Liu et al. (2022). Both chemicals are of great industrial interest (i.e., pharmaceutical, feed, chemical, biofuels). Optimization of AF processes towards the production of either ethanol or iso-butyric acid or both can be an additional approach for cheese whey valorization.

T_PT_9 NP and T_PT_7 NP displayed a production of acetic acid of 1,479.13 and 1,558.17 mg COD_AcOH_ L^-1^, respectively. As shown in figure 2, there was no production of methane. Acetic acid was consumed in A_PT_9NP, whereas in A_PT_7NP its production was slightly lower than both T_PT_9 NP and T_PT_7 NP with 1,056.04 mg COD_AcOH_ L^-1^.

Interestingly, CNP also presented a higher acetic acid production of 1,109.13 mg COD_AcOH_ L^-1^, which shows that headspace pressure is a more suitable selective pressure parameter than inoculum pre-treatment for acetic acid production. The produced acetic acid can thus be utilized as an organic carbon source for higher added-value biological processes (i.e., microalgal photoorganoheterotrophic biomass production).

The imposed parameters (*i.e*., inoculum pre-treatments, initial pH, and headspace pressure) also acted as selective pressures in the batch experiments modifying the microbial fermentation end-products. In general, thermal pre-treatment, non-pressurized headspace, and initial pH 9.0 displayed a trend for higher VFA production. Nevertheless, it is not possible to affirm what the best combination of parameters would be since there was not a pattern of production observed for all VFAs in this given setup.

However, the combination of parameters can be suitable for the production of specific volatile acids or ethanol, as seen in T_PT_7NP and T_PT_9NP for the production of acetic (1,479.13 and 1,558.17 mg COD_AcOH_ L^-1^, respectively) and butyric acids (1,207.46 and 1,425.36 mg COD_BTA_ L^-1^), and A_PT_9NP for iso-butyric acid and ethanol (7,677.30 mg COD_IBA_ L^-1^ and 13,683.24 mg COD_EtOH_ L^-1^).

Non-pressurized batches presented a global production of 3 times more VFAs than pressurized batches. Besides common VFAs, pressurized headspace batches produced formic and lactic acids but in negligible quantities as shown in table 3.

Initial WPC40 carbohydrate concentration was approximately 260 mg_glucose_ L^-1^, stabilizing at 65 mg_glucose_ L^-1^ on the fourth day. The acidogenesis rate of 1.9 d^-1^, was close to the typical rate of 2 d^-1^ observed in similar studies(van Lier et al., 2008). COD removal and volatile solids (VS) showed similar patterns, with sCOD decreasing from 4.2 to 2.4 g L^-1^ and VS decreasing from 4.92 to 4.74 g L^-1^. This was likely due to biomass production and organic matter oxidation. Control batches showed a decrease from 4.92 to 4.74 g L^-1^, corroborating with COD removal results. Cheese whey alkalinity varied by 6% in all batches (1.53 to 1.45 CaCO_3_ L^-1^). Acidification of whey might thus have led to the production of VFAs, as seen in both control and thermal pre-treatment pH 9.0. Table 4 shows the results of the main physicochemical analyses.

**Table 4.** Initial and final results of the main physicochemical analyses performed in the AF experiment for 30 days. Batches consisted of control, alkaline and thermal pre-treatments with initial pH 7.0 and 9.0 (A_PT_7, A_PT_9, T_PT_7, and T_PT_9) with pressurized (P) and non-pressurized (NP) headspaces. Analyses considered are: chemical oxygen demand (COD), total organic carbon (TOC), total nitrogen (TN), total solids (TS), total volatile solids S), and pH.

| Batch operational conditions |  |  |  |  |  |  |  |  |  |  |  |
| --- | --- | --- | --- | --- | --- | --- | --- | --- | --- | --- | --- |
| Analyses (mg L<sup>-1</sup>)<br>* × • | Time | Control |  | APT7 |  | APT9 |  | TPT7 |  | TPT9 |  |
|  |  | P | NP | P | NP | P | NP | P | NP | P | NP |
| CH<sub>4</sub>• | t0 | 0.00 | 0.00 | 0.00 | 0.00 | 0.00 | 0.00 | 0.00 | 0.00 | 0.00 | 0.00 |
|  | tf | 11,086.37 | 5.69 | 0.00 | 0.54 | 52.29 | 0.67 | 0.00 | 0.00 | 0.00 | 0.00 |
| COD* | t0 | 8,368.63 | 8,634.33 | 7,336.90 | 11,174.50 | 8,583.00 | 15,294.50 | 9,325.30 | 8,758.00 | 10,000.00 | 9,736.00 |
|  | tf | 7,391.10 | 11,053.99 | 7,694.78 | 10,658.81 | 7,610.05 | 10,584.72 | 8,859.74 | 14,289.56 | 8,779.31 | 13,844.98 |
| TOC | t0 | 2,005.67 | 1,695.33 | 3,280.00 | 1,326.00 | 2,890.00 | 1,636.00 | 3,467.00 | 2,230.00 | 3,691.00 | 2,237.00 |
|  | tf | 2,284.00 | 1,181.00 | 2,229.00 | 1,213.00 | 2,385.00 | 1,176.00 | 3,133.00 | 1,842.00 | 3,266.00 | 1,784.00 |
| TN | t0 | 333.40 | 617.99 | 309.80 | 69.25 | 306.50 | 100.30 | 468.30 | 219.20 | 463.50 | 216.10 |
|  | tf | 279.33 | 174.97 | 268.20 | 157.40 | 403.60 | 136.80 | 544.90 | 317.60 | 484.10 | 284.20 |
| C: N ratio | t0 | 6.02:1 | 2.7:1 | 10.59:1 | 19.15:1 | 9.43:1 | 16.31:1 | 7.40:1 | 10.17:1 | 7.96:1 | 10.35:1 |
|  | tf | 8.18:1 | 6.75:1 | 8.31:1 | 7.71:1 | 5.91:1 | 8.60:1 | 5.75:1 | 5.80:1 | 6.75:1 | 6.28:1 |
| TS | t0 | 0.00516 | 5.72 | 0.00478 | 6.69 | 0.00539 | 6.71 | 0.00729 | 7.68 | 0.00698 | 8.52 |
|  | tf | 0.00323 | 5.50 | 0.00344 | 5.08 | 0.00330 | 4.69 | 0.00376 | 6.13 | 0.00401 | 5.64 |
| TVS | t0 | 0.00453 | 4.98 | 0.00421 | 5.87 | 0.00468 | 5.84 | 0.00578 | 6.81 | 0.00600 | 7.40 |
|  | tf | 0.00264 | 4.66 | 0.00278 | 4.23 | 0.00260 | 3.99 | 0.00311 | 5.10 | 0.00324 | 4.58 |
| pH | t0 | 8.25 | 8.25 | 7.00 | 7.00 | 9.00 | 9.00 | 7.00 | 7.00 | 9.00 | 9.00 |
|  | tf | 4.76 | 5.14 | 4.65 | 4.99 | 5.00 | 5.01 | 4.87 | 5.11 | 5.06 | 5.19 |

#### 3.1.3 Inoculum thermal pre-treatment, initial pH, and non-pressurized headspace acted as selective pressure mechanisms on microbial communities during cheese whey acidogenic fermentation

In the replicate experiment performed in TU Delft, both control and inoculum thermal pre-treated microbial communities’ composition and evolution were determined through 16S rRNA gene amplicon sequencing (Figures 3 and 4). Initial microbial communities were highly similar, with few quantitative and only minor qualitative differences being observed. They both shared 7 of their 8 main detected phyla, 10 of their 11 main detected classes and 18 of their 19 main detected orders, as well as their 10 and 5 most important detected families and genera, respectively

**Figure 3.**
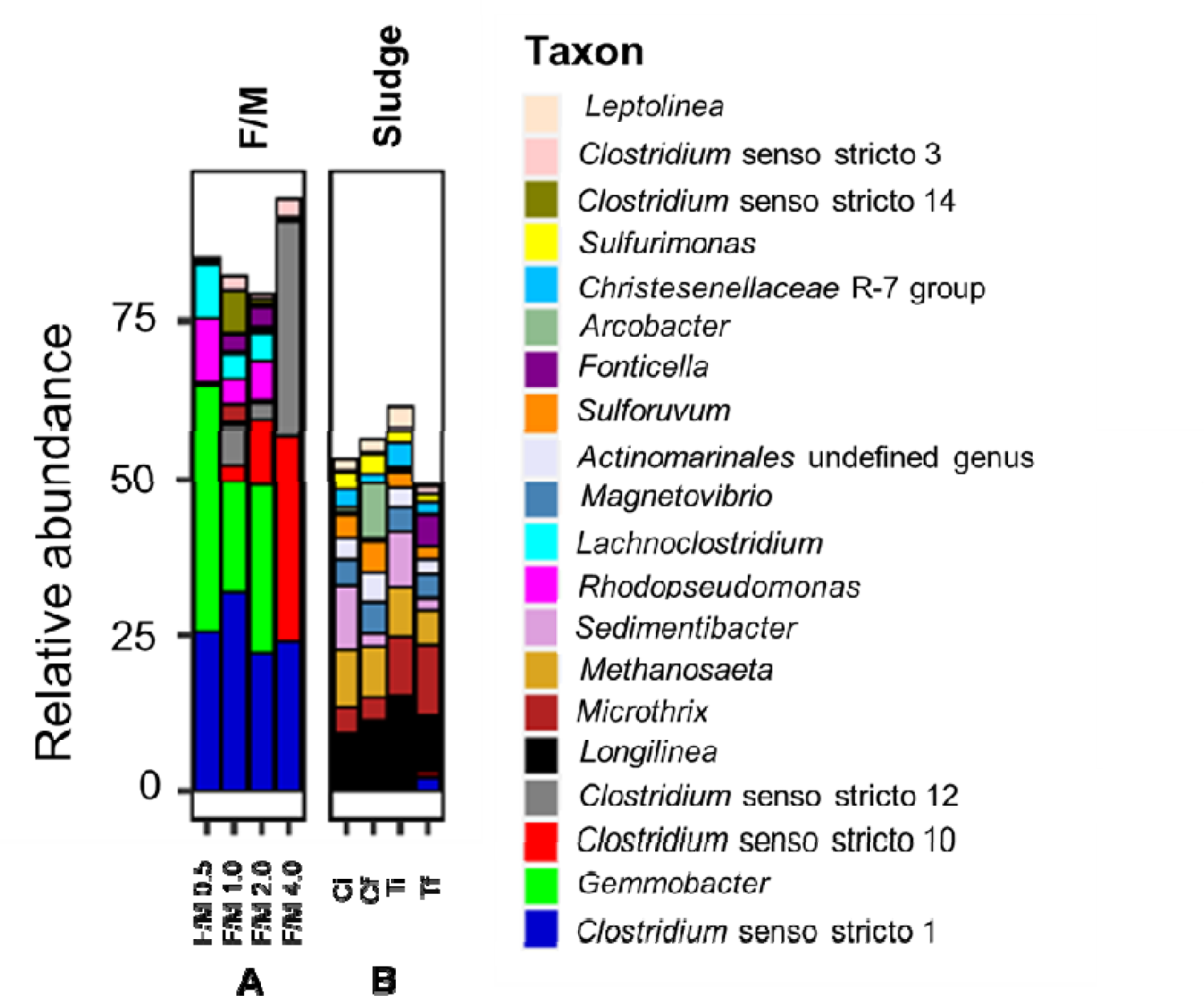
Microbial communities’ compositions and evolution. The 20 most abundant archaeal and bacterial genera identified through 16S rRNA gene amplicon sequencing. A microbial population from the F/M ratios experiments (FM). B, microbial populations from the control and thermal pre-treated inoculum batches (T_PT_9 NP).

**Figure 4.**
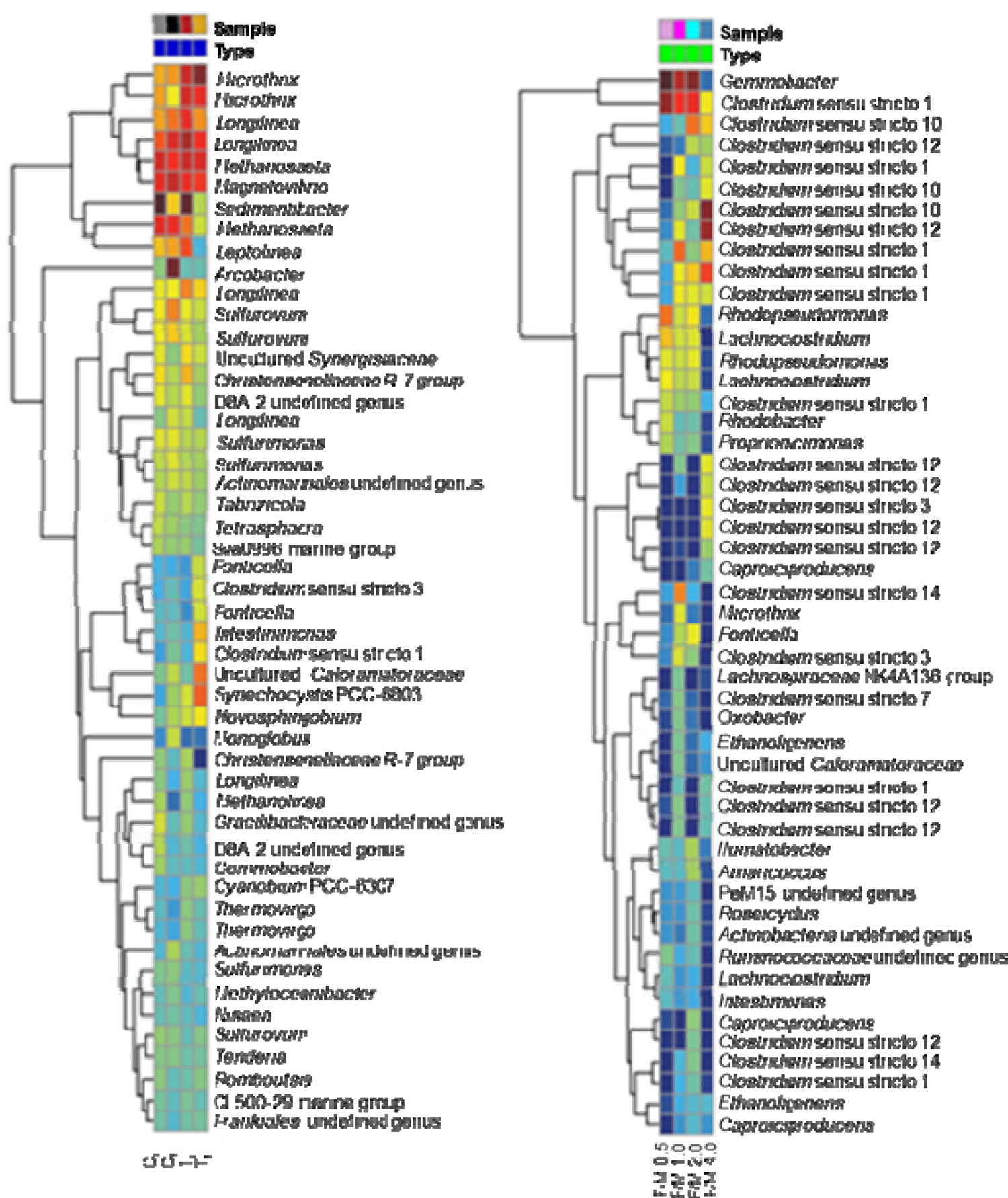
ASV abundance. The 50 most abundant archaeal and bacterial ASVs detected by the AmpliconTagger pipeline. FM: F/M ratios experiments; T_PT_NP9: thermal pre-treated inoculum batches.

As shown in figures 3 and 4, the common genera included fermenters and VFAs producers (such as *Sedimentibacter* (Lin et al., 2019), *Longilinea*, *Leptolinea* (Dai et al., 2016), and *Christensenellaceae* R-7 group (Y. Gao et al., 2022), the lipid-accumulating and bulk and foaming *Candidatus Microthrix* (Sheik et al., 2016), microorganisms involved in sulfur and nitrogen metabolisms (*Magnetovibrio* (Bazylinski et al., 2013), *Sulfurovum* and *Sulfurimonas* (Li et al., 2018), as well as the acetoclastic methanogen *Methanosaeta* (Stams et al., 2019).

The only noticeable difference consisted of *Cyanobacteria* being detected in the inoculum thermal pre-treated population (Figure 4). Cyanobacteria are aerobic photosynthetic microorganisms (Stal and Moezelaar, 1997). However, they display mechanisms that enable them to survive in unfavourable conditions such as in hypoxic or anaerobic (Li et al., 2018) environments. Although chlorophyll (Chl) deficiency can be caused by low levels of oxygen, these microorganisms can regulate Chl production by activating oxygen-independent oxidases or by inducing the transcription of genes that encodes enzymes that work in microoxic conditions (Fujita et al., 2015). In the given scenario, facultative anoxygenic photosynthesis uses sulfide as the electron donor with photosystem I-driven photoassimilation (Hamilton et al., 2018).

Control and inoculum thermal pre-treated microbial communities evolved differently along the process. The final populations presented highly similar compositions at the phylum, class, order, and family levels, sharing all or almost all the main detected taxons with only quantitative differences being perceived. Noticeable qualitative and quantitative differences were however observed at the genus level (Figures 3 and 4). Figure 5 depicts the beta diversity analysis of the samples.

**Figure 5.**
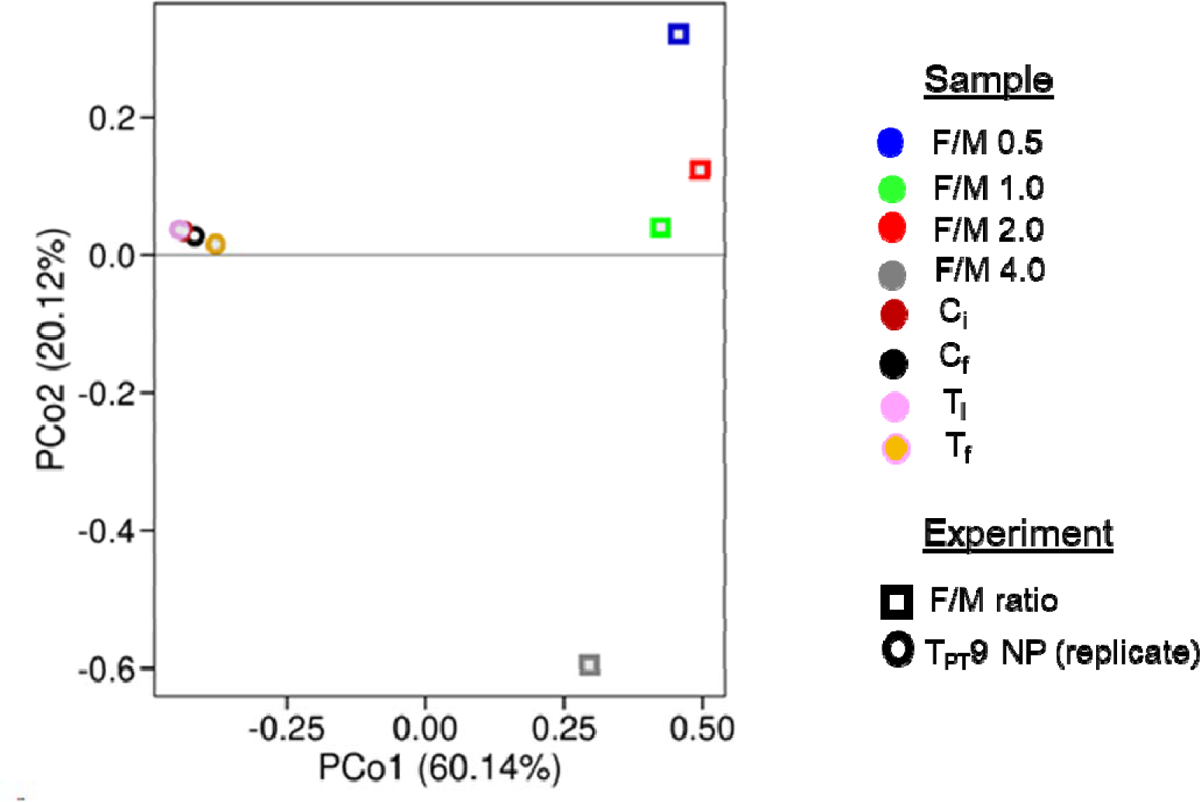
Beta diversity analysis of the microbial populations. Principal Coordinates Analysis computed from Bray-Curtis dissimilarity distance matrix. FM: F/M ratios experiments; T_PT_NP9: thermal pre-treated inoculum batches.

In the control final microbial community, an increase was observed in fermenters, such as *Longilinea* and *Leptolinea* (Dai et al., 2016), and in microorganisms involved in sulfur and nitrogen metabolisms, such as *Magnetovibrio* (Bazylinski et al., 2013), *Sulfurovum* and *Sulfurimonas* (Li et al., 2018). *Arcobacter* (S. Gao et al., 2022), a sulfide oxidizer and denitrifier, also appeared and became the second genus of importance, while the two VFAs producers *Sedimentibacter* (Lin et al., 2019) and *Christensenelaceae* R-7 group (Y. Gao et al., 2022) strongly decreased.

In opposition, in the inoculum thermal pre-treated final microbial community, an increase was observed in VFAs and alcohols producers, such as *Clostridium* (Mockaitis et al., 2022), *Fonticella* (Joshi et al., 2021), and *Intestinimonas* (Kang et al., 2021), while fermenters such as *Longilinea* and *Leptolinea* strongly decreased. The lipid-accumulating and bulk and foaming *Candidatus Microthrix* (Sheik et al., 2016) and the *Cyanobacteria Synechocystis* (Fujita et al., 2015) increased, and *Magnetovibrio* (Bazylinski et al., 2013), *Sulfurovum* and *Sulfurimonas* (Li et al., 2018), involved in sulfur and nitrogen metabolisms, slightly decreased.

Those results could explain the differences in performance observed in both processes. It is likely that methanogenesis was halted in the inoculum thermal pre-treated assay by the strong decrease observed in microorganisms such as *Longilinea* and *Leptolinea*, which are fermenters known to provide substrates to methanogens (Yamada et al., 2007, 2006), concomitantly with the strong increase in VFAs and alcohols producers.

The decrease, even if limited, observed in *Methanosaeta* tends to confirm that methanogens were inhibited. The presence of *Methanosaeta* could be explained by several hypotheses: (*i*) samples detected were dead biomass or spores, (*ii*) inoculum pre-treatments were sufficient to inhibit or diverge the metabolic pathway for methane formation, or (*iii*) they could have been outcompeted by acetate fermenters (Callbeck et al., 2019; Miñana-Galbis et al., 2002; Nuppunen-Puputti et al., 2018; Shi et al., 2020). Further-omics studies can clarify this finding. The presence of denitrifiers could be a response to batches oxygen limitation since they are aerobic facultative (Skiba, 2008). They can also compete for nitrate content in cheese whey (Oliveira et al., 1995).

Initial WPC40 carbohydrate concentration was approximately 260 mg_glucose_ L^-1^, stabilizing at 65 mg_glucose_ L^-1^ on the fourth day. The acidogenesis rate of 1.9 d^-1^, was close to the typical rate of 2 d^-1^ observed in similar studies(van Lier et al., 2008). COD removal and volatile solids (VS) showed similar patterns, with sCOD decreasing from 4.2 to 2.4 g L^-1^ and VS decreasing from 4.92 to 4.74 g L^-1^. This was likely due to biomass production and organic matter oxidation. Control batches showed a decrease from 4.92 to 4.74 g L^-1^, corroborating with COD removal results. Cheese whey alkalinity varied by 6% in all batches (1.53 to 1.45 CaCO_3_ L^-1^). Acidification of whey might thus have led to the production of VFAs, as seen in both control and thermal pre-treatment pH 9.0.

### 3.2 Impact of F/M ratio on VFAs production during acidogenic fermentation of cheese whey

#### 3.2.1 Lower F/M ratio is more efficient for VFAs production

Four different F/M ratios of WPC40 were evaluated in an experiment performed with a thermal pre-treated inoculum at initial pH of 9.0 and under non-pressurized conditions. Ratios were F/M 0.5, F/M 1.0, F/M 2.0, and F/M 4.0, corresponding to 0.5, 1.0, 2.0, and 4.0 g COD g VSS^-1^, respectively.

The F/M 0.5 ratio acted as a positive control since the thermal batch in the replicate experiment presented the same F/M ratio of 0.5 g COD g VSS^-1^. About 95% of carbohydrates were consumed by the third day, while substrate acidification started in the first couple of days F/M 0.5 (1,056 mg COD_compound_ L^-1^ d^-1^), F/M 1.0 (789 mg COD_compound_ L^-1^ d^-1^), F/M 0.2 (486 mg COD_compound_ L^-1^ d^-^1nd F/M 4.0 (390 mg COD_compound_ L^-1^ d^-1^).

As shown in Table 6, F/M 0.5 ratio showed the best VFA production (3,317.88 mg COD_compound_ L^-1^), whereas the F/M ratio 2.0 (345.93 mg L^-1^) had the lowest. Non-pressurized thermal pre-treatment batch in the replicate experiment had a VFAs production of 2,568.26 mg L^-1^. There was no production of methane in all samples which corroboratto the results observed in previous experiments.

**Table 5.** VFAs spectra of replicated control and thermal pre-treatment (pH 9.0) acidogenic fermentation experiment compared to the original experiment for 10 days. Replicate results are expressed as a means average of the triplicate assays. (--) The product was not evaluated.

| VFAs Production (mg COD compound L<sup>-1</sup>) |  |  |  |  |  |
| --- | --- | --- | --- | --- | --- |
| Products | Time | UNICAMP |  | TU Delft |  |
|  |  | C | T | C | T |
| Acetic acid | t0 | 154.34 | 308.26 | 0.00 | 19.28 |
|  | tf | 1,037.34 | 1,361.07 | 319.13 | 1,407.90 |
| Butyric acid | t0 | 0.00 | 0.00 | 0.00 | 0.00 |
|  | tf | 187.29 | 1121.49 | 5.35 | 106.91 |
| Iso-butyric acid | t0 | 0.83 | 0.00 | 16.04 | 149.67 |
|  | tf | 0.00 | 0.00 | 16.04 | 58.80 |
| Caproic acid | t0 | 0.00 | 0.00 | 290.94 | 179.70 |
|  | tf | 0.00 | 0.00 | 479.20 | 2,587.25 |
| Iso-caproic acid | t0 | -- | -- | 0.00 | 0.00 |
|  | tf | -- | -- | 0.00 | 42.79 |
| Ethanol | t0 | 0.00 | 0.00 | -- | -- |
|  | tf | 0.00 | 155.07 | -- | -- |
| Formic acid | t0 | 0.00 | 0.00 | 4.30 | 4.30 |
|  | tf | 0.00 | 0.00 | 0 | 0.00 |
| Lactic acid | t0 | 0.00 | 1,137.59 | -- | -- |
|  | tf | 0.00 | 0.00 | -- | -- |
| Malic acid | t0 | 0.00 | 24.74 | -- | -- |
|  | tf | 0.00 | 0.00 | -- | -- |
| Propionic acid | t0 | 0.00 | 0.00 | 11.19 | 11.19 |
|  | tf | 161.48 | 55.21 | 85.76 | 55.93 |
| Valeric acid | t0 | 0.00 | 0.00 | 0.00 | 6.94 |
|  | tf | 0.00 | 0.00 | 20.83 | 27.78 |
| Iso-valeric acid | t0 | 0.00 | 0.00 | 0.00 | 20.83 |
|  | tf | 46.54 | 235.89 | 20.83 | 83.34 |

**Table 6.** VFAs spectra of F/M ratios experiments. F/M 0.5, F/M 1.0, F/M 2.0, and F/M 4.0: 0.5, 1.0, 2.0 and 4.0 g COD g VSS^-1^, respectively. F/M 0.5 acted as a positive control as it has the same 0.5 g COD g VSS^-1^ ratio as the replicated thermal pre-treatment batch in section 3.2. The experiment was performed in triplicate and lasted 10 days.

| VFAs Production (mg CODcompound L<sup>-1</sup>) |  |  |  |  |  |
| --- | --- | --- | --- | --- | --- |
| Products | Time | F/M 0.5 | F/M 1.0 | F/M 2.0 | F/M 4.0 |
| Acetic acid | t0 | 34.02 | 9.19 | 0.00 | 0.00 |
|  | tf | 900.48 | 956.51 | 493.42 | 865.84 |
| Butyric acid | t0 | 0.00 | 0.00 | 0.00 | 0.00 |
|  | tf | 349.75 | 25.26 | 34.64 | 36.48 |
| Iso-butyric acid | t0 | 1,707.03 | 78.98 | 60.70 | 25.58 |
|  | tf | 43.22 | 38.89 | 19.32 | 12.99 |
| Caproic acid | t0 | 154.28 | 30.42 | 0.00 | 0.00 |
|  | tf | 1,823.41 | 1,042.24 | 3,620.88 | 194.20 |
| Iso-caproic acid | t0 | 0.00 | 0.00 | 0.00 | 0.00 |
|  | tf | 0.00 | 0.00 | 0.00 | 0.00 |
| Formic acid | t0 | 1.85 | 2.21 | 2.89 | 3.33 |
|  | tf | 0.00 | 0.00 | 0.30 | 0.00 |
| Propionic acid | t0 | 5.26 | 0.00 | 0.00 | 0.00 |
|  | tf | 66.33 | 34.40 | 7.27 | 23.27 |
| Valeric acid | t0 | 9.79 | 0.00 | 0.00 | 0.00 |
|  | tf | 45.21 | 0.00 | 27.19 | 4.90 |
| Iso-valeric acid | t0 | 32.61 | 17.61 | 0.00 | 0.00 |
|  | tf | 89.48 | 58.23 | 40.31 | 27.40 |
| Total production |  | 3,317.88 | 2,155.53 | 4,211.46 | 1,165.08 |

#### 3.2.2 F/M ratios affected microbial communities’ evolution

Microbial communities evolved differently during the F/M experiments. As shown in Figure 5, increasing F/M ratios significantly impacted the microbial populations’ evolution, with the strongest impact observed at the highest ratio of 4.0. Under such conditions, the microbial population was ultra-dominated by a single genus, *Clostridium* (Figures 3 and 4). At lower F/M ratios, the community was mainly composed of the same four genera, which altogether represented 84.3%, 77.6%, and 74.7% of the F/M-0.5, F/M-1, and F/M-2 populations, respectively (Figure 3 and 4). Those four genera were *Gemmobacter*, a poly-hydroxybutyrate-accumulating and denitrifying bactéria (Wang et al., 2019), *Clostridium*, a known fermenter and VFAs producer (Mockaitis et al., 2022), *Lachnoclostridium*, a known butyrate producer (Detman et al., 2021), and *Rhodopseudomonas*, a highly metabolically versatile bacteria capable of hydrogen production through nitrogen fixation or poly-hydroxybutyrate production (Ranaivoarisoa et al., 2019). The operational conditions applied in the F/M-0.5 experiment were thus apparently those that favor the best microbial combination to produce the highest level of VFAs. Further characterizations (i.e., metagenomics and/or metatranscriptomics analyses) have to be performed to better understand why such close microbial populations could generate different amounts of VFAs.

## 4 Conclusions

AF is a successful alternative for producing VFAs of economic interest out of high-strength residues such as cheese whey. Inoculum pre-treatments can play as selective pressures on microbial communities, selecting microorganisms that can thrive in imposed conditions. The main conclusions from this work are the following:

1. Thermal pre-treatment of inocula was efficient to halt methanogenesis regardless of headspace pressure, pH, and F/M ratios.
2. Contrary to the literature, alkaline pre-treatment did not improve methanogenesis. However, non-pressurized headspace alkaline pre-treatment pH 9.0 did produce significant quantities of iso-butyric acid (7,677.30 mg COD_IBA_ L^-1^) and ethanol (13,693.24 mg COD_EtOH_ L^-1^). Further studies on the mechanisms of this pre-treatment are necessary to optimize the process.
3. *Methanosaeta* was present in the replicate of the control (pH 8.25) and thermal (pH 9.0) batches, despite the absence of methane production. Thermal pre-treatment could have either inactivated or destroyed this microorganism, which in its turn enabled the detection of *Methanosaeta* in the 16S rRNA gene amplicon sequencing analysis.
4. A low F/M ratio of 0.5 g COD g^-1^ VS selected a microbial community producing a high level of VFAs by AF.
5. The setup of parameters used as selective pressure during cheese whey AF must vary according to the desired end product.
6. Headspace pressure directs the metabolic pathways for VFAs and alcohol production in AF. Pressurized headspace batches had a more di while non-pressurized batches produced higher amounts of VFAs. The exception would be iso-butyric acid and ethanol produced in non-pressurized alkaline pH 9.0 batches.
7. Another important parameter was the Initial pH of 9.0. Regardless of the headspace pressure and inoculum pre-treatment, an initial pH of 9.0 produced the greatest number of products. The drastic pH drop influences the redox potential of the medium facilitating the uptake of compounds while setting the grounds for ecologic relationships within the microbial community.

This work aimed to verify if imposed parameters (i.e., headspace pressure, inoculum thermal and alkaline pre-treatment, and initial pH) in cheese whey AF would work as selective pressures on the inoculum microbial community. Our objectives were met when we managed to halt methanogenesis and increase acetic acid, iso-butyric acid, and ethanol. Acetate is an organic carbon source that can be used for different biological conversions (e.g., microalgal photoorganoheterotrophic processes), whereas iso-butyric and ethanol are of interest of pharmaceutical, feed, chemical, and biofuels industries.

Still, there is plenty to be understood, especially regarding the influence of these parameters on metabolic pathways and microbial interaction. The fact that thermal pre-treatment time and temperature were optimized and methanogenesis was halted, despite the presence of methanogens. Understanding the mechanism behind this can overcome competition for substrate.

Thermal inoculum pre-treatment can be used in short fermentation processes. Nevertheless, the interaction among the factors with different substrates, inoculum and reactor configurations must be studied to certify thermal pre-treatment efficiency. Thus, it is necessary to evaluate other configurations for process implementation and optimization.

Understanding the VFAs production and consumption mechanisms is crucial for driving the spectrum of fermentation products to desired compounds. Future 16S rRNA metagenomics analyses will be fundamental to decrease these knowledge gaps.

## Acknowledgements

This work was funded in major CAPES PDS scholarship (CAPES PDS 88882.435082/2019-01) and CNPq (CNPq 166460/2017-6). The work at the TU Delft was funded by the start-up grant of the Department of Biotechnology of the TU Delft (Prof. Dr. David Weissbrodt, PI). The lab works at UNICAMP benefited from the assistance of Vítor Augusto de Oliveira, Juliana Martins Valença, Giovani Archanjo Brotto and Rosa Helena Aguiar while the lab works in the TU Delft benefited from the assistance of Cor Ras, Johan Knoll, Marcel Langerveld, Ben Abbas and Armand Middeldorp.

## 6 Authors’ contributions

M.P.G. d A conceptualized the experiment and wrote the manuscript with direct core inputs by D.G.W and G.M. M.P.G.dA performed the analysis for the acidogenic fermentation in Brazil and in The Netherlands, the latter with the assistance of C.M. The analysis and conceptualisation the F/M experiments were performed by C.M. The experiment was designed by MPGdA, DGW and GM by confronting ideas, concepts and solutions to technological, outcomes on microbial communities genomics were provided by G.B and J.T. All authors read, edited, and provided critical feedback to the manuscript.

